# Connectomic analysis of taste circuits in *Drosophila*

**DOI:** 10.1101/2024.09.14.613080

**Authors:** Sydney R. Walker, Marco Peña-Garcia, Anita V. Devineni

**Author notes:** These authors contributed equally (Sydney R. Walker, Marco Peña-Garcia).

## Abstract

Our sense of taste is critical for regulating food consumption. The fruit fly *Drosophila* represents a highly tractable model to investigate mechanisms of taste processing, but taste circuits beyond sensory neurons are largely unidentified. Here, we use a whole-brain connectome to investigate the organization of *Drosophila* taste circuits. We trace pathways from four populations of sensory neurons that detect different taste modalities and project to the subesophageal zone (SEZ). We find that second-order taste neurons are primarily located within the SEZ and largely segregated by taste modality, whereas third-order neurons have more projections outside the SEZ and more overlap between modalities. Taste projections out of the SEZ innervate regions implicated in feeding, olfactory processing, and learning. We characterize interconnections between taste pathways, identify modality-dependent differences in taste neuron properties, and use computational simulations to relate connectivity to predicted activity. These studies provide insight into the architecture of *Drosophila* taste circuits.

## INTRODUCTION

Our sense of taste is critical in helping us determine what foods to eat. Attractive tastes indicate that food contains calories or nutrients, while aversive tastes warn that the food may be toxic or spoiled. Most animals, including humans, recognize five basic tastes: sweet, umami (savory), salty, bitter, and sour. In general, taste sensory cells express receptors for individual taste modalities and thus respond to specific tastes.^1^ In mammals, signals from taste sensory cells in the tongue are transmitted to the brainstem, thalamus, and gustatory cortex.^2^ Neural recordings have revealed a variety of response types across these areas, including cells that respond to specific tastes (“specialists”), cells that respond broadly to multiple tastes (“generalists”), and cells with varying response dynamics.^3^ Despite this progress, it is not clear how these different types of neuronal responses arise from the connectivity of the circuit or how specific response types contribute to behavior. Addressing these questions requires a system where we can examine the connectivity, response properties, and behavioral contribution of individual cell types, thus determining how these properties are related.

The fruit fly *Drosophila melanogaster* represents such a system. Recent studies have generated a brain-wide map of neuronal connectivity^4^ and classified the ∼130,000 brain neurons into thousands of cell types,^5^ each typically consisting of just one or a few cells that can be genetically targeted using collections of transgenic driver lines.^6–11^ Taste cells in *Drosophila* are distributed throughout multiple organs, including the proboscis (the feeding organ), legs, and wings.^12^ The major taste organ that regulates feeding is the labellum, located at the distal end of the proboscis. Several classes of gustatory receptor neurons (GRNs) have been identified in the labellum, including sugar-sensing, bitter-sensing, water-sensing, high salt-sensing, and IR94e-expressing neurons.^12–14^ Sugar GRNs promote feeding, whereas bitter and high salt GRNs suppress feeding.^13,15^ Water-sensing GRNs detect solutions of low osmolarity and promote water consumption.^16^ IR94e GRNs respond to low salt concentrations and amino acids, and these GRNs suppress feeding while promoting egg-laying.^13,17,18^

In contrast to our understanding of taste coding at the periphery, downstream taste circuits in the *Drosophila* brain have remained largely unknown. Labellar GRNs project axons into the subesophageal zone (SEZ) of the brain,^19,20^ but their postsynaptic partners remained a mystery for decades. The development of trans-Tango, a trans-synaptic tracing method, enabled the visualization of neurons that receive synaptic input from GRNs, termed second-order taste neurons.^21^ Tracing the postsynaptic partners of sugar or bitter GRNs revealed a large population of second-order neurons with extensive projections in the SEZ and a few major projections to higher brain regions.^21–23^ Second-order sugar and bitter neuron populations are anatomically similar and appear to include some overlapping neurons.^22^ However, the large number of neurons labeled by trans-Tango made it difficult to determine the extent of overlap and, more generally, to identify individual types of second-order neurons.

The recent release of the first *Drosophila* whole-brain connectome^4^ makes it possible to identify the inputs and outputs of every neuron, enabling us to trace the flow of taste information through the brain. Two initial studies (using connectome data prior to public release) identified labellar GRNs in the connectome dataset^24^ and traced a circuit connecting sugar GRNs to a motor neuron that drives proboscis extension, which represents the initiation of feeding.^25^ The sugar proboscis extension circuit consists of three layers of interneurons located within the SEZ.^25^ An additional study simulated whole-brain activity using a connectome-based model and predicted which neurons are activated by each taste modality, finding substantial overlap between the neurons activated by sugar and water GRNs but not other modalities.^17^

Several fundamental questions regarding the architecture of *Drosophila* taste circuits remain unanswered. First, how much taste processing occurs within the SEZ, and which higher brain regions receive input from each taste modality? Second, how much overlap exists between taste circuits for different modalities, and does this overlap increase or decrease at subsequent layers of processing? Third, to what extent do taste circuits consist of feedforward excitation, feedforward inhibition, or lateral and feedback connections, and does this depend on the taste modality?

In this study, we addressed these questions by analyzing the whole-brain connectome released by the FlyWire consortium,^4,5^ which also contains neurotransmitter predictions for each neuron.^26^ We first traced the postsynaptic partners of GRNs to identify second-order taste neurons. We found that second-order neurons are primarily located within the SEZ, are largely segregated by modality, and show modality-dependent differences in the location and neurotransmitter type of their output projections. Third-order taste neurons showed more overlap between modalities and more extensive projections outside the SEZ, primarily innervating brain regions in the superior protocerebrum in modality-specific patterns. Finally, we used simulations of whole-brain activity to analyze the relationship between connectivity and predicted activity. Together, these studies provide insight into the architecture of circuits for taste processing in the fly brain, laying the groundwork for functional studies.

## RESULTS

### Second-order taste neurons are largely modality-specific

We started by tracing the postsynaptic partners of labellar GRNs, focusing on the four major GRN classes that have been previously annotated in the connectome: sugar-sensing, bitter-sensing, water-sensing, and IR94e-expressing neurons. We identified GRNs in the connectome based on GRN lists from recent studies,^17,18^ neuron annotations in FlyWire, and manual inspection of each neuron’s morphology (see Methods). We limited our analyses to GRNs on the left side of the labellum, which project to the left hemisphere of the brain, because these neurons have been more thoroughly annotated and we are more confident of their identity. Our GRN lists include 22 sugar GRNs, 18 water GRNs, 20 bitter GRNs, and 9 IR94e-expressing GRNs (Figure 1A; note that neuron images from FlyWire are always left-right inverted). Sugar and water GRNs project to overlapping regions of the SEZ, whereas the projections of bitter and IR94e GRNs are more distinct. GRNs are thought to be excitatory, which aligns with most of the neurotransmitter predictions in FlyWire^26^ and experimental data showing that second-order sugar neurons are activated by sugar.^25^

**Figure 1.**
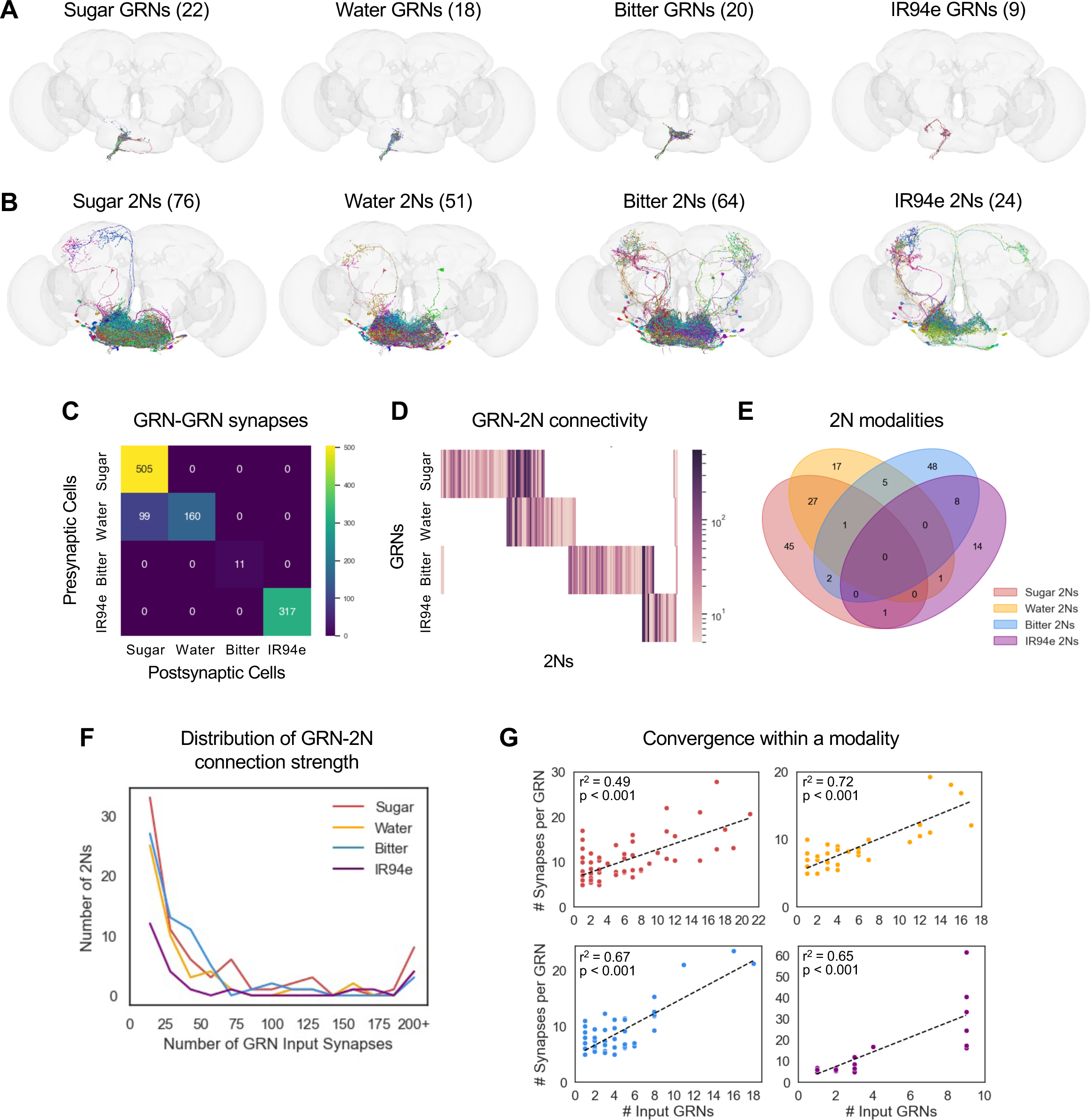
2Ns are largely specific to individual taste modalities. (A-B) Images of GRNs (A) and 2Ns (B) from each of the four taste modalities. Numbers in parentheses denote the number of neurons in each category. Note that neuron images from FlyWire are always left-right inverted. (C) Heatmap showing the number of GRN-GRN synapses for each pair of modalities. (D) Heatmap showing GRN-2N connectivity for each individual 2N. The colorbar represents the total number of input synapses from each GRN type. (E) Venn diagram showing overlap between 2Ns of different modalities. Classification of 2Ns by local vs. projection

We then used the connectome to identify second-order taste neurons (2Ns), neurons that receive direct input from GRNs. Using a connection threshold of 5 synapses, a threshold suggested by FlyWire in order to exclude false positives,^4^ we identified 76 sugar 2Ns, 51 water 2Ns, 64 bitter 2Ns, and 24 IR94e 2Ns (Figure 1B). These numbers correspond to approximately three times as many 2Ns as GRNs, with 2N/GRN ratios ranging from 2.7 for IR94e to 3.5 for sugar.

We also identified many GRN-GRN connections, which have been described in a previous study.^24^ The majority of these connections were between GRNs of the same type, with the exception of water GRNs synapsing onto sugar GRNs (Figure 1C). When normalized by the number of GRNs, within-type connections were most prominent for IR94e (35 synapses/GRN), followed by sugar GRNs (23 synapses/GRN) and water GRNs (9 synapses/GRN), and they were almost nonexistent for bitter GRNs (< 1 synapse/GRN). Thus, some taste modalities exhibit more crosstalk between GRNs than others. Note that GRNs were excluded from our lists of 2Ns, as our goal was to trace feedforward taste pathways.

The 2Ns for each taste modality were largely distinct, with the exception of strong overlap between sugar/water 2Ns and weaker overlap between bitter/IR94e 2Ns and bitter/water 2Ns (Figure 1D-E). For example, of the 76 sugar 2Ns, 28 also receive input from water GRNs, whereas only 3 receive input from bitter GRNs and only 1 receives input from IR94e GRNs.

Within a modality, the strength of GRN input to each 2N varied widely, with the majority of 2Ns receiving less than 10 GRN synapses while other 2Ns received over 100 synapses (Figure 1F). 2Ns received a median of 15-17 total synapses from a median of 2-3 GRN input cells, depending on modality. Interestingly, 2Ns receiving input from more GRNs also tended to receive more synapses per GRN input cell, whereas fewer input GRNs corresponded with weaker connections (Figure 1G; r^2^ values ranged from 0.49 to 0.72). This correlation would tend to widen the distribution of GRN-2N connection strength, creating a larger separation between 2Ns receiving few versus many GRN inputs. Together, these analyses show that 2Ns are largely segregated by modality – with notable exceptions for the convergence of sugar/water and bitter/IR94e inputs – and vary widely in the strength of the taste input they receive.

### Properties of 2Ns vary by taste modality

We next examined the anatomical and predicted functional properties of 2Ns. We classified 2Ns as local neurons, whose outputs reside exclusively within the SEZ, or projection neurons, which have at least some outputs outside of the SEZ. The majority of 2Ns were local neurons, and the number of projection neurons varied by modality: projection neurons comprised only 8-10% of sugar or water 2Ns (5-6 cells) but comprised 31% of bitter 2Ns (20 cells) and 54% of IR94e 2Ns (13 cells) (Figure 2A-B). Consistent with these numbers, the majority of 2N output synapses were located within the SEZ, and the proportion of output synapses outside of the SEZ was much higher for bitter and IR94e 2Ns (21%) than water and sugar 2Ns (3-6%) (Figure 2C).

**Figure 2.**
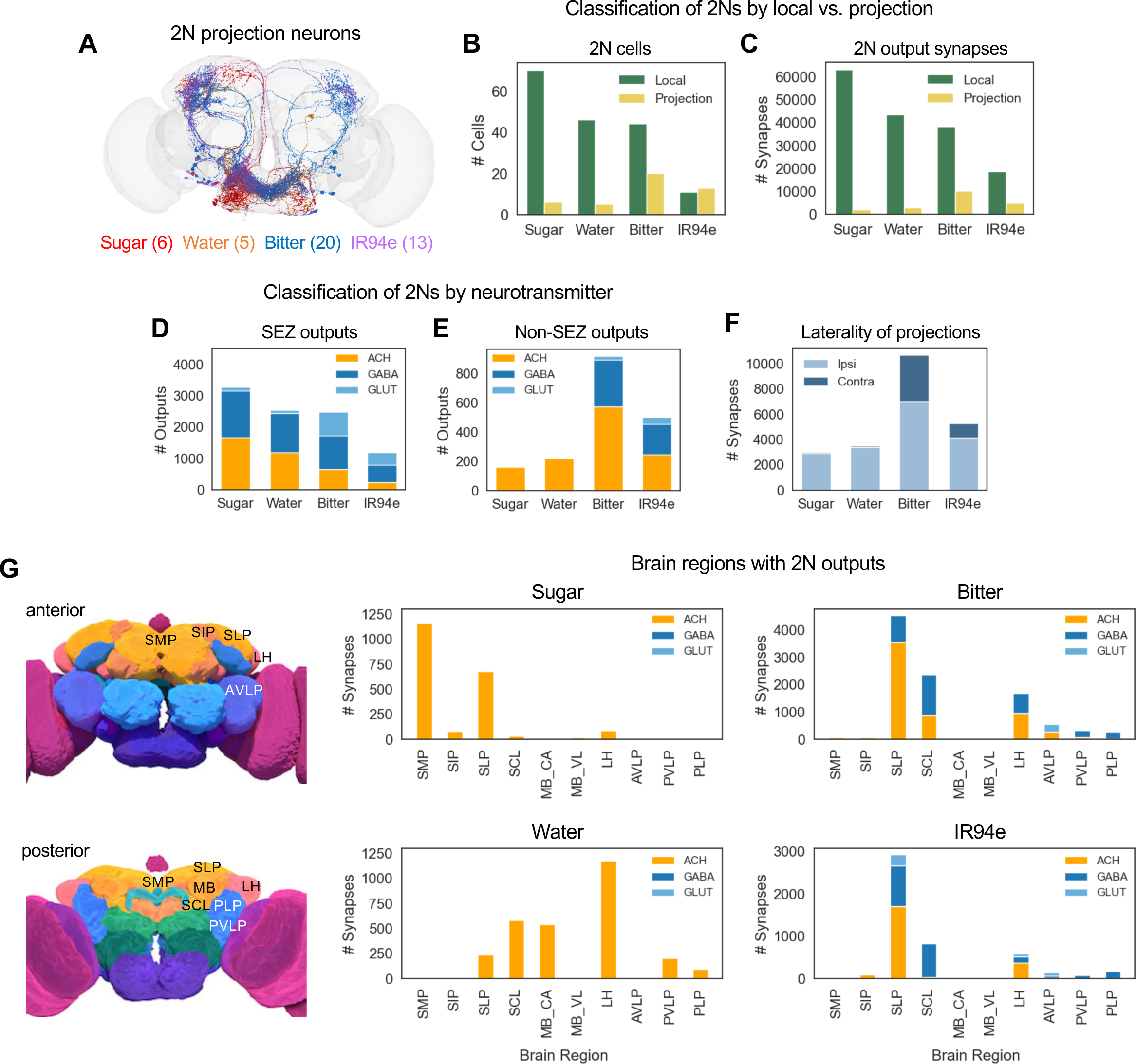
Anatomical and functional properties of 2Ns vary by taste modality. (A) Image showing 2N projection neurons for each modality. Neurons are shown at 50% opacity to visualize neurons at multiple depths and enable intermediate coloring for neurons belonging to more than one modality, although these represented a very small proportion of projection neurons. (B-C) Proportion of 2Ns (B) and 2N output synapses (C) that are local to the SEZ versus projecting out of the SEZ. (B) All pairwise comparisons of the proportion of local vs. projection neurons between modalities were significantly different except for sugar vs. water and bitter vs. IR94e (Fisher’s exact test). (C) All pairwise comparisons of the proportion of local vs. projection output synapses between modalities were significantly different except for bitter vs. IR94e (Fisher’s exact test). (D-E) Predicted neurotransmitters for 2N local (D) and projection (E) output connections. All pairwise comparisons of the proportion of excitatory vs. inhibitory neurons (D) or outputs (E) between modalities were significantly different except for sugar vs. water (Fisher’s exact test). (F) Proportion of 2N output synapses outside of the SEZ that are located in the ipsilateral versus contralateral hemisphere (relative to the location of GRN projections). All pairwise comparisons of the proportion of ipsilateral vs. contralateral outputs between modalities were significantly different except for sugar vs. water (Fisher’s exact test).

We then classified 2Ns by their predicted neurotransmitter type.^26^ In the *Drosophila* central nervous system, the major excitatory neurotransmitter is acetylcholine and the major inhibitory neurotransmitters are GABA and glutamate.^27,28^ We compared the predicted neurotransmitters used by 2Ns projecting within and outside the SEZ. 2N outputs within the SEZ included both excitatory and inhibitory connections, with the ratio of these types being roughly equal for sugar and water 2Ns (49-53% inhibitory) and skewed toward inhibition for bitter (73% inhibitory) and IR94e (81% inhibitory) 2Ns (Figure 2D). Interestingly, 2N outputs outside of the SEZ had a different distribution that was much more skewed toward excitation rather than inhibition: they were only 2% inhibitory for sugar and water 2Ns and 38% or 51% inhibitory for bitter and IR94e 2Ns, respectively (Figure 2E). Thus, we observe differences in the predicted neurotransmitter types for 2Ns that depend on their taste modality and where they project.

We next analyzed the anatomical locations of 2N projections outside the SEZ. The majority of 2N output synapses were located in the ipsilateral hemisphere of the brain (relative to the location of GRN projections), but bitter and IR94e 2Ns had a much higher proportion of contralateral outputs (34% and 22%, respectively) than sugar and water 2Ns (3%) (Figure 2F). Projection 2Ns had output synapses in a variety of brain regions, and the location of outputs varied by modality (Figure 2G). Bitter and IR94e projection 2Ns, which make up the majority of projection 2Ns, had outputs predominantly in the superior lateral protocerebrum (SLP), superior clamp (SCL), and lateral horn (LH). Sugar projection 2Ns had outputs primarily in the SLP and the superior medial protocerebrum (SMP), and water projection 2Ns had outputs in the LH, SCL, SLP, and mushroom body calyx. Together, these analyses reveal that 2Ns primarily convey taste information within the SEZ but also project to a small number of higher-order brain regions in a modality-specific manner.

### Third-order taste neurons overlap across modalities

Next, we identified the postsynaptic partners of 2Ns. 2Ns had a substantial number of connections with each other, including connections between all possible pairs of modalities (Figure 3A). The most common connections were within the same modality or between sugar and water 2Ns, similar to GRN-GRN connections. We separately analyzed only excitatory or inhibitory synapses and found that cross-modal connections were biased towards being inhibitory rather than excitatory (Figure 3B-C). 2Ns also had many synapses onto GRNs, and cross-modal 2N-GRN connections were most prominent between the sugar and water pathways and between the bitter and IR94e pathways (Figure 3D), similar to the pattern of GRN-2N convergence (Figure 1E).

**Figure 3.**
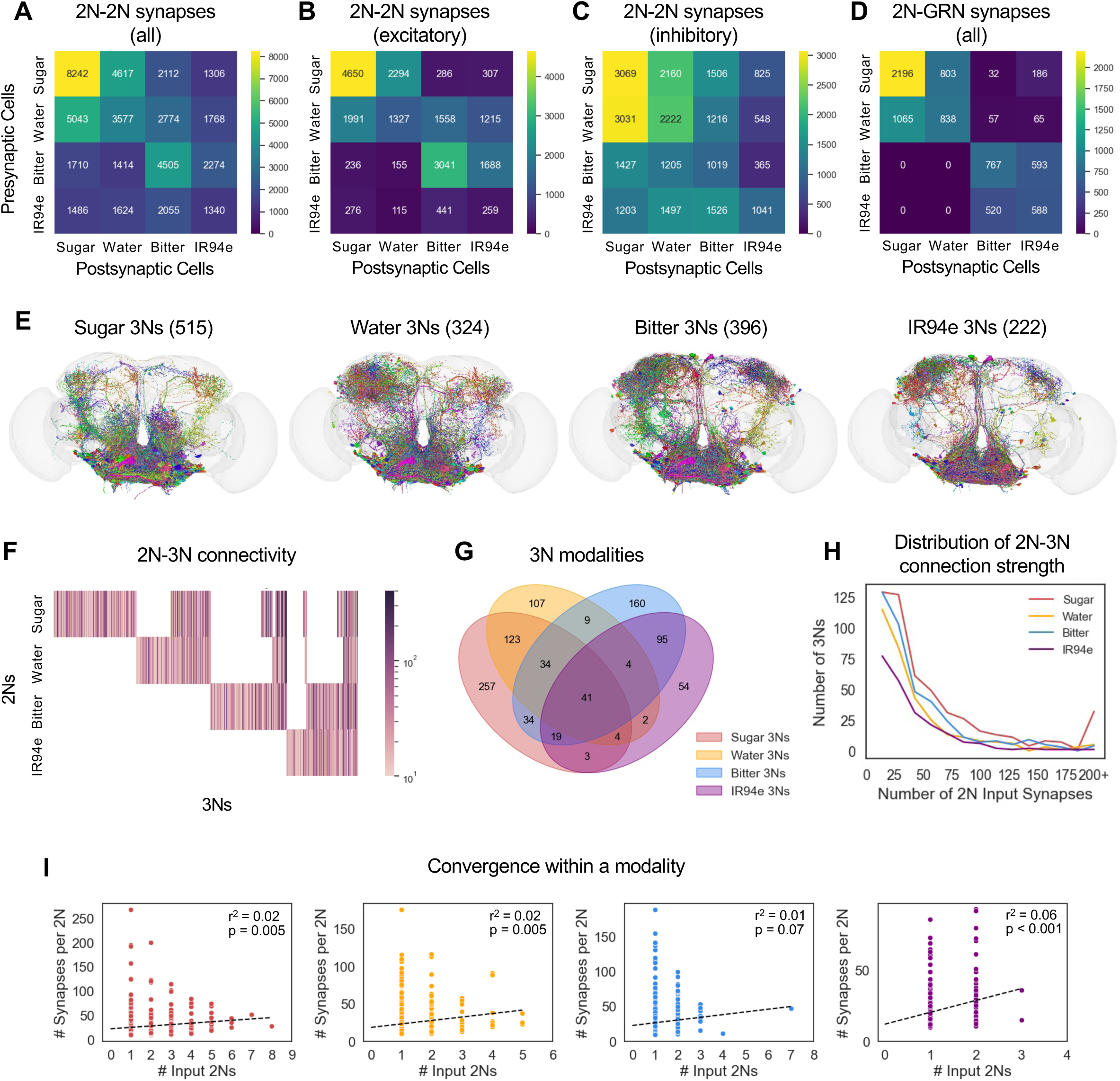
3Ns overlap across taste modalities. (A-D) Heatmaps showing the number of 2N output synapses onto other 2Ns (A-C) or to GRNs (D). Panel A shows all 2N-2N synapses, whereas panels B and C show only excitatory or inhibitory synapses, respectively. (E) Images of 3Ns for each of the four taste modalities. Numbers in parentheses denote the number of 3Ns. (F) Heatmap showing 2N-3N connectivity for each 3N. The colorbar represents the total number of input synapses from each 2N type. (G) Venn diagram showing overlap between 3Ns of different modalities. (H) Distribution of 2N-3N connection strength (number of total 2N input synapses to each 3N) for each modality.

We then focused on 2N outputs to third-order neurons (3Ns). Because iterative tracing of connectivity generates larger and larger numbers of neurons at each layer, we set a more stringent threshold for 3N identification: we only traced the partners of 2Ns receiving at least 10 total synapses from GRNs of the same modality, and we similarly set a 10 synapse cutoff for 2N to 3N connections within a modality. Our list of 3Ns excludes GRNs of any type as well as neurons that are 2Ns for the same modality, but a 3N could be a 2N for a different modality. With these criteria, we identified 515 sugar 3Ns, 324 water 3Ns, 396 bitter 3Ns, and 222 IR94e 3Ns (Figure 3E).

3Ns showed much more overlap between taste modalities than 2Ns, with some overlap between every possible pair of modalities (Figure 3F-G). For example, of the 515 sugar 3Ns, 39% were water 3Ns, 25% were bitter 3Ns, and 13% were IR94e 3Ns. Overall, 50-76% of 3Ns received input from more than one modality (50% of sugar 3Ns, 67% of water 3Ns, 60% of bitter 3Ns, and 76% of IR94e 3Ns).

Within a modality, 3Ns received a median of 22-29 total synapses (depending on modality) from a median of one 2N (Figure 3H), representing a lower rate of convergence than observed from GRNs to 2Ns (but note that the criteria for 2N-3N connectivity were more stringent). Unlike GRN to 2N synapses, for 2N to 3N synapses there was no clear correlation between the number of input cells and the strength of each input connection (Figure 3I; r^2^ values ranged from 0.01-0.06).

We analyzed whether 2Ns from the same modality tend to provide input of the same sign onto 3Ns. Convergent input of the same sign would lead to net excitation or inhibition, whereas convergent input of opposite signs would be expected to cancel out. Focusing on 3Ns that receive convergent input from two or three 2Ns of the same modality, which represent 86% of all 3Ns receiving multiple 2N inputs, we found that the frequency of convergent input of the same sign (all excitatory or all inhibitory) was always higher than the frequency expected by chance, especially for excitatory convergence (Figure 4A-B).

**Figure 4.**
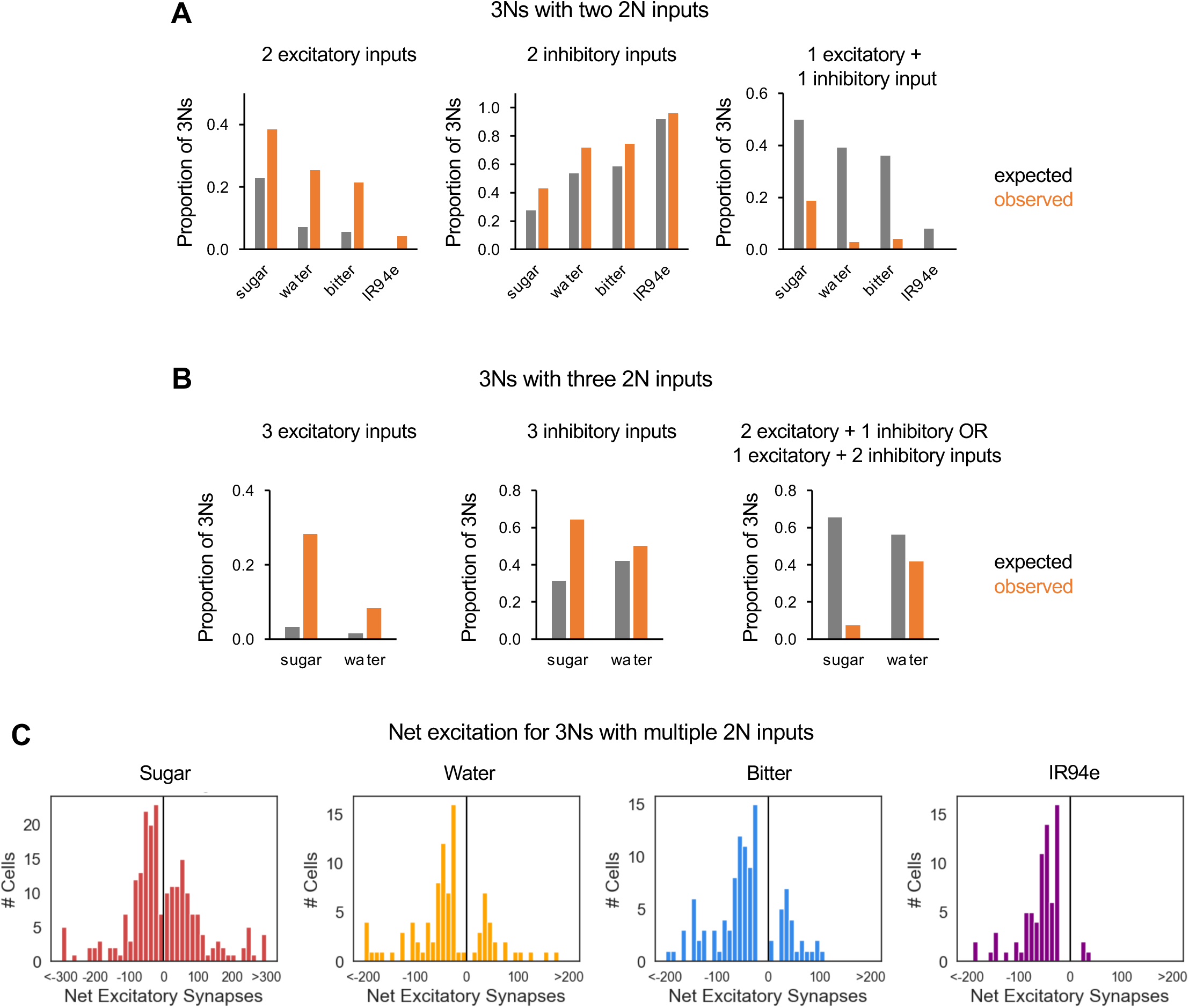
2N inputs of the same sign tend to converge onto 3Ns. (A) 3Ns receiving exactly two 2N inputs from the same modality were analyzed. The expected proportion of 3Ns receiving two excitatory inputs (left), two inhibitory inputs (middle), or one excitatory and one inhibitory input (right) was compared to actual proportions. The observed proportion of 3Ns in each category differed significantly from expected proportions for each of the four modalities (p<0.001, chi-squared test; n = 71-112 3Ns depending on modality). (B) 3Ns receiving exactly three 2N inputs from the same modality were analyzed. The expected proportion of 3Ns receiving three excitatory inputs (left), three inhibitory inputs (middle), or a combination of excitatory and inhibitory inputs (right) was compared to actual proportions. We only included modalities for which we could analyze at least 10 3Ns (sugar and water). The observed proportion of 3Ns in each category differed significantly for sugar (p<0.001, chi-squared test; n = 53 3Ns) but not for water, which had a small sample size (n = 12 3Ns). (C) Distribution of net excitation onto each 3N that receives multiple 2N inputs within a modality. Net excitation represents the difference between the number of excitatory and inhibitory 2N input synapses; positive numbers represent net excitation and negative numbers represent net inhibition.

We also calculated the net excitation onto each 3N receiving multiple 2N inputs by quantifying the difference in the total number of excitatory and inhibitory input synapses from 2Ns. 3Ns ranged from receiving strong net excitation to strong net inhibition, with relatively few neurons near zero (Figure 4C), consistent with the observation that inputs of the same sign tend to converge (Figure 4A-B). 3Ns of all modalities other than sugar appeared to be skewed toward receiving net inhibition rather than net excitation, suggesting that many 3Ns are inhibited rather than activated by taste stimuli. Together, these results show that different taste pathways begin to converge at the level of 3Ns, with inputs from the same modality expected to produce net excitation or inhibition.

### 3Ns expand taste processing to additional brain regions

To further characterize the 3Ns, we classified them based on anatomical and functional types. As with the 2Ns, we first classified 3Ns based on where their output synapses reside. A small proportion of 3Ns did not have any output synapses within the brain (6% of sugar 3Ns and 1-2% of 3Ns for other modalities), and these neurons are primarily motor neurons or descending neurons that project to the ventral nerve cord. Excluding these neurons, we classified 3Ns based on whether their output synapses reside exclusively within the SEZ (local neurons) or whether some outputs reside outside the SEZ (non-local neurons, which include neurons projecting out of the SEZ as well as neurons residing entirely in other brain regions) (Figure 5A-C). Similar to 2Ns, the majority of 3Ns were local SEZ neurons (Figure 5C). However, the proportion of non-local sugar and water 3Ns (17% or 35%, respectively) was substantially higher than the proportion of non-local sugar and water 2Ns (8% or 10%, respectively).

**Figure 5.**
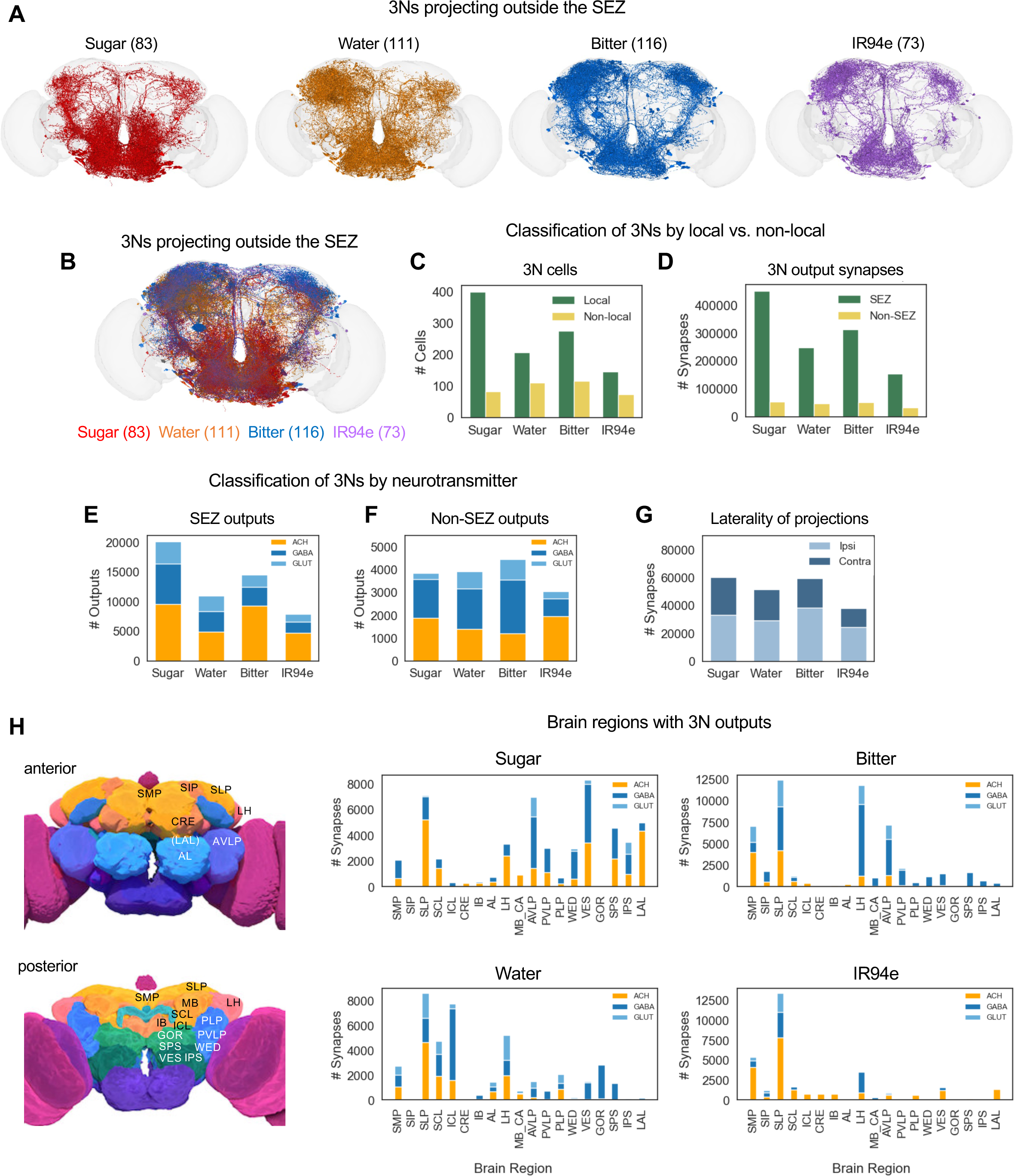
3Ns expand taste processing to additional brain regions. (A-B) Images showing 3Ns of each modality that have output synapses outside of the SEZ. 3Ns for individual modalities (A) are shown in addition to the overlaid image (B) to facilitate comparisons of 3N projection patterns across modalities, given the high density of projections and the substantial overlap between 3Ns of different modalities. Neurons in panel B are shown at 50% opacity. (C-D) Proportion of 3Ns (B) and 3N output synapses (C) that are that are local to the SEZ versus outside of the SEZ. (B) Pairwise comparisons of the proportion of local vs. non-local neurons between modalities revealed a significant difference between sugar and the other three modalities but not between any other modalities (Fisher’s exact test). (C) All pairwise comparisons of the proportion of SEZ vs. non-SEZ outputs between modalities were significantly different (Fisher’s exact test). (E-F) Predicted neurotransmitters for 3N output connections within the SEZ (D) or outside of the SEZ (E). All pairwise comparisons of the proportion of excitatory vs. inhibitory neurons (D) or outputs (E) between modalities were significantly different (Fisher’s exact test). (G) Proportion of 3N output synapses outside of the SEZ that are located in the ipsilateral versus contralateral hemisphere (relative to the location of GRN projections). All pairwise comparisons of the proportion of ipsilateral vs. contralateral outputs between modalities were significantly different except for sugar vs. IR94e (Fisher’s exact test). (H) Number of 3N output synapses in brain regions outside of the SEZ, color-coded by neurotransmitter type. The top 12 brain regions for each modality are included, comprising a total of 19 regions that include >97% of non-SEZ output synapses for sugar, bitter, and IR94e 3Ns and 91% for water 3Ns. Pictures on the left show the approximate location of each brain region (images are from the FlyWire website). Brain regions in parentheses are not visible in the plane shown but are located in a slightly more posterior plane. See text and Methods for abbreviations.

Proportions of non-local bitter and IR94e 3Ns were also high (30% or 33%, respectively) but not higher than the proportion of non-local 2Ns for these modalities (31% or 54%, respectively). Consistent with the preponderance of local 3Ns, a large majority of 3N output synapses were located within the SEZ (82-89% for all modalities; Figure 5D). Thus, while the third layer of the taste pathway involves an expansion to brain regions beyond the SEZ, taste processing within the SEZ continues to dominate.

We next classified 3Ns by their predicted neurotransmitter type. For the sugar and water modalities, 3N outputs within the SEZ were roughly equally distributed between excitatory and inhibitory connections (45-48% excitatory; Figure 5E), similar to sugar and water 2Ns. Interestingly SEZ outputs from bitter and IR94e 3Ns were skewed toward excitation (61-64% excitatory; Figure 5E), in contrast to SEZ outputs from bitter and IR94e 2Ns that were skewed toward inhibition (see Figure 2D). Some modalities showed a difference in the proportion of excitatory versus inhibitory 3N outputs when comparing SEZ and non-SEZ synapses (Figures 5E-F); for example, bitter 3N outputs within the SEZ were much more likely to be excitatory than those outside the SEZ. Thus, like 2Ns, the predicted neurotransmitter type of 3Ns depends on their modality and location.

We then analyzed where 3Ns send output projections outside of the SEZ. Although 3Ns had more ipsilateral than contralateral output synapses (relative to the location of GRN projections), similar to 2Ns, the proportion of contralateral outputs was much higher for 3Ns (35-44%) than for 2Ns (3-34%) (Figure 5G). 3Ns had strong outputs in some of the same areas innervated by 2Ns, including the SMP, SLP, and LH, but 3Ns also projected to additional areas, such as the anterior ventrolateral protocerebrum (AVLP) (Figure 5H). The location of 3N projections varied substantially by modality (Figure 5H). For example, sugar 3Ns had many more outputs in the lateral accessory lobe (LAL) and ventromedial neuropils (vest, VES; superior posterior slope, SPS; inferior posterior slope, IPS) than other types of 3Ns. Together, these results show that 3Ns convey taste information both within and outside the SEZ with modality-specific patterns.

### Relating connectivity to simulated neuronal activity

A recent study by Shiu et al. developed a computational model to simulate neuronal activity across the fly brain based on each neuron’s connectivity and predicted neurotransmitter.^17^ Simulations rely on a leaky integrate-and-fire model, and the parameters of the model (e.g., membrane properties, resting potential, action potential threshold) were fit with experimental data. Although this model has limitations, such as assuming that each neuron has the same biophysical properties, Shiu et al. found that the model was remarkably accurate in predicting activity within sensory-motor circuits, including the sugar taste circuit.^17^ We used this model to ask whether the taste 2Ns and 3Ns that we identified are predicted to respond to taste stimulation and what factors determine their predicted response.

First, we asked how many of the 2Ns that we identified are activated by GRN stimulation in the model. We stimulated GRNs of each taste modality at intensities ranging from 25 to 200 Hz. GRN stimulation generally elicited activity in dozens to hundreds of neurons, corresponding to less than 1% of all neurons in the brain, with the exception of IR94e stimulation at the highest intensities (Figure 6A). In contrast, the proportion of 2Ns activated by GRN stimulation was 1-2 orders of magnitude higher, ranging from 10-60% at lower intensities to 70-80% at higher intensities (Figure 6B). Across all stimulation intensities, 2Ns receiving a larger number of synaptic inputs from GRNs were more likely to be activated (Figure 6C). The median number of GRN input synapses was typically >40 for activated 2Ns and less than 10 for non-activated 2Ns, with the disparity being largest at low stimulation intensities. These results suggest that although the majority of 2Ns are expected to be activated by taste input, a substantial proportion of 2Ns may receive GRN input that is too weak to elicit neuronal firing on its own, potentially contributing to neuronal activity in combination with other inputs.

**Figure 6.**
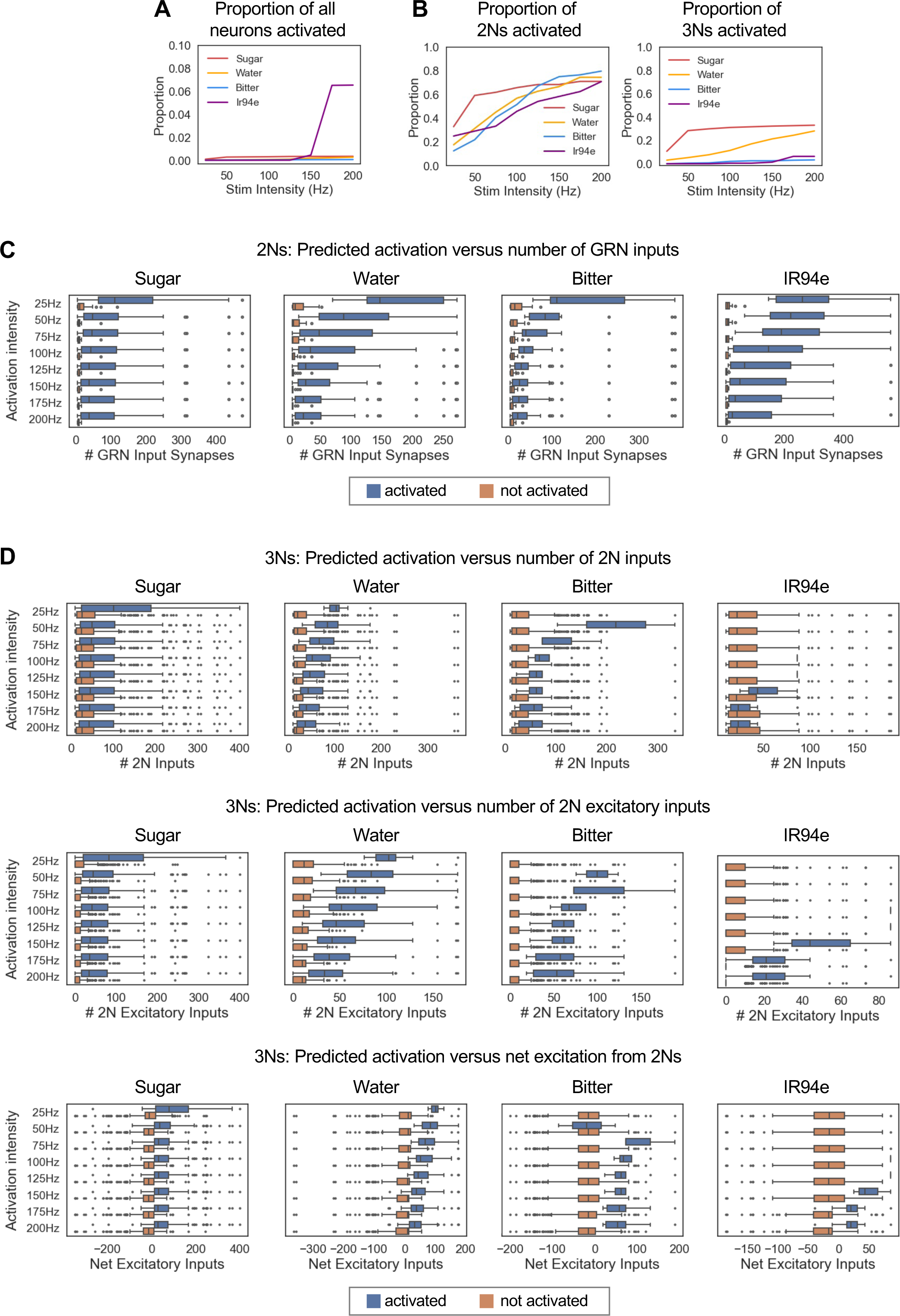
Relating connectivity to simulated neuronal activity. (A) Proportion of all neurons in the brain that were activated by simulated GRN activation at each intensity. Note the y-axis scale is different from the scale in panel B. Less than 1% of neurons were activated under most conditions, with the exception of IR94e stimulation at the highest intensities, which activated ∼6% of neurons. (B) Proportion of 2Ns (left) or 3Ns (right) for each modality that were activated by simulated GRN activation at each intensity. (C) Comparison of the number of GRN input synapses for 2Ns that were activated vs. not activated in the simulations. (D) Comparison of the number of 2N input synapses (top row), 2N excitatory input synapses (middle row), or net excitatory input synapses (the difference between the number of 2N excitatory and inhibitory input synapses; bottom row) for 3Ns that were activated vs. not activated in the simulations. In panels C and D, boxes represent the interquartile range with a line at the median, bars represent 1.5 times the interquartile range, and points represent outliers outside of this range.

We then asked how many of the 3Ns are activated by GRN stimulation in the model. For each modality, the proportion of activated 3Ns was much lower than the proportion of activated 2Ns, ranging from 0-11% at lower stimulation intensities to 3-33% at higher intensities (Figure 6B). A much higher proportion of sugar and water 3Ns were predicted to be activated than bitter and IR94e 3Ns (28-33% for sugar and water versus 3-6% for bitter and IR94e at the highest intensity). Similar to 2Ns, 3Ns receiving a larger number of synaptic inputs from 2Ns were more likely to be activated, but the difference in input between activated and non-activated neurons was not as striking as for 2Ns (Figure 6D, top). We hypothesized that this may be because 2N inputs to 3Ns can be either excitatory or inhibitory; excitatory 2Ns would promote 3N activation while inhibitory 2Ns would suppress 3N activation. Consistent with this hypothesis, activated versus non-activated 3Ns showed a clearer difference in their number of excitatory 2N inputs (Figure 6D, middle) or their net excitation from 2Ns (Figure 6D, bottom) as compared to the number of total 2N inputs (Figure 6D, top). The median number of excitatory 2N synapses was ∼40-100 for activated 3Ns and ∼0-10 for non-activated 3Ns, roughly similar to the numbers for 2Ns. Together, these results show that a lower proportion of 3Ns than 2Ns are predicted to be activated by a given taste input, and this may reflect the use of inhibitory taste coding at this layer as well as the need for excitatory summation from multiple stimuli beyond a single taste input.

## DISCUSSION

In this study, we used the whole-brain connectome to analyze taste pathways in the central brain. We found that 2Ns are largely segregated by modality, with the notable exception of overlap between sugar and water 2Ns. 2Ns primarily convey taste information within the SEZ but also project to a small number of higher-order brain regions in a modality-specific manner. Compared to 2Ns, 3Ns showed more overlap between modalities and more extensive projections outside the SEZ. 3Ns received both excitatory and inhibitory inputs from 2Ns, with input of the same type tending to converge within a modality. Connectome-based simulations revealed that the majority of 2Ns, but only a minority of 3Ns, are predicted to be activated by GRN stimulation, suggesting that some 2Ns and 3Ns may require additional input for activation or, in the case of 3Ns, may encode taste through an inhibition of activity. Together, these studies provide an overview of early taste processing in *Drosophila* that will guide future work aimed at studying specific neurons within these circuits.

### Tracing central taste circuits

For decades, central taste circuits in *Drosophila* have remained largely unknown. Initially the field relied on functional or anatomical screens to identify neurons involved in taste processing, revealing interneurons and motor neurons within the SEZ.^29–31^ However, pan-neuronal imaging suggested that taste is processed in both the SEZ and the higher brain,^32^ consistent with studies showing that taste input is relayed to the mushroom body to mediate taste learning.^33,34^ The first systematic approach to identify central taste pathways emerged with the development of trans-Tango,^21^ revealing that second-order sugar and bitter neurons comprise large populations that extensively innervate the SEZ with a few projections to the superior protocerebrum.^21–23^

Our analysis of the connectome provides a much more comprehensive and high-resolution view of taste processing at both the second and third layers. Our analyses of 2Ns are largely consistent with results of previous studies, finding that 2Ns reside primarily within the SEZ with selective projections to other brain regions. However, the 2Ns identified from the connectome do not appear to include all of the projection neurons labeled with trans-Tango.^21–23^ Some neurons labeled with trans-Tango but not identified by connectomic analyses may receive taste input from neurons outside of the labellum, such as the legs or pharynx, as our study only traced input from labellar GRNs whereas Gal4 drivers for trans-Tango studies label GRNs in multiple organs. It is also possible that some neurons identified by trans-Tango represent false positives, as trans-Tango signals are based on proximity whereas connectome data is based on electron microscopy, a much more reliable method to identify synapses.

### Overlap between taste modalities

The extent of overlap between central pathways for processing different taste modalities has been a longstanding subject of debate in both insects and mammals.^3,12^ Imaging studies in *Drosophila* suggested that sweet and bitter pathways are largely non-overlapping,^32^ but recordings in other insects identified central neurons activated by multiple taste modalities.^35,36^ Our analysis of the connectome reveals that 2Ns are largely segregated by modality, with the exception of strong overlap between sugar/water and weaker overlap between bitter/IR94e and bitter/water. The extent of overlap likely relates to the behavioral role of each taste modality: both sugar and water promote ingestive behaviors, whereas bitter strongly suppresses feeding and IR94e has a modest inhibitory role in feeding.^12,13,15–18^

Our finding that sugar and water pathways show strong convergence aligns with the results of Shiu et al., who used whole-brain simulations to analyze the overlap between neurons activated by each GRN type,^17^ as well as a recent imaging study by Li et al. that examined taste coding across cell bodies in the SEZ.^37^ In addition to finding strong overlap between sugar- and water-responsive neurons, Li et al. found that taste-responsive SEZ cells (which may represent 2Ns, 3Ns, or neurons farther downstream) showed a variety of tuning profiles, with approximately one-third of cells responding to multiple tastes.^37^ These findings are consistent with our results showing a strong convergence of inputs from different taste modalities by the third layer. Our analyses revealed more convergence across modalities (50-76% of 3Ns receive input from multiple modalities) than the proportion of multimodal cells identified by Li et al., but convergence in our study includes both excitatory and inhibitory inputs. We found that cross-modal inhibition between 2Ns was more common than cross-modal excitation, and this inhibition would serve to sparsen gustatory tuning rather than broaden it. Together, these new analyses and imaging studies suggest that the fly taste system may contain a diversity of response types, similar to the case in mammals.^3^ This diversity of tuning may be important for taste discrimination or for the activation of circuits that mediate different behaviors.

### Brain regions for taste processing

Proboscis motor neurons that control feeding behaviors are located in the SEZ,^29,38^ suggesting that taste circuits do not need to project out of the SEZ to regulate innate feeding programs.

Indeed, a recent study identified a five-layered circuit contained entirely within the SEZ that drives proboscis extension to sugar,^25^ and SEZ circuits also mediate locomotor stopping in response to sweet taste.^39,40^ However, other behaviors influenced by taste, such as egg-laying,^41,42^ associative learning,^33,43,44^ or multisensory integration may require taste information to be transmitted outside of the SEZ. Our results show that although early taste processing occurs primarily within the SEZ, some neurons relay information to higher brain regions and the extent of non-SEZ processing expands at the third layer. At the second layer, more bitter and IR94e 2Ns project out of the SEZ as compared to sugar and water 2Ns. This difference may reflect specific behavioral programs driven by bitter or IR94e neurons; for example, IR94e 2Ns projecting to the SLP have been implicated in the regulation of egg-laying.^18^

We found that major targets of taste input include the SLP, SMP, and LH. The LH is a well-known olfactory region, as it receives input from second-order olfactory neurons and mediates innate responses to odors.^45^ The LH has strong connections to and from the SLP,^4^ a neighboring region, suggesting that odor and taste inputs may be integrated in these two areas. The SMP contains neurosecretory cells that regulate feeding behaviors, including insulin-producing cells,^46^ and SMP neurons regulate sugar and water consumption.^47,48^ Interestingly, the projections of 2Ns and 3Ns vary substantially based on taste modality. For example, only sugar 2Ns project to the SMP, only water 2Ns project to the mushroom body calyx, and sugar 3Ns have much stronger projections to the LAL and ventromedial neuropils (VES, SPS, IPS) than other 3N types. These differences suggest that certain modalities have more important roles in regulating behaviors linked to those brain regions, such as feeding (SMP) or learning (mushroom body), providing hypotheses for future studies to investigate.

We also observed differences in the balance of excitatory and inhibitory connections based on their anatomical location and modality. Local 2Ns in the bitter and IR94e pathways were more likely to be inhibitory than sugar and water 2Ns, which may reflect the need for bitter and IR94e neurons to inhibit feeding circuits in the SEZ. 2N outputs were more likely to be excitatory if they were located outside of the SEZ than within the SEZ, suggesting that projection neurons relay long-range excitation to circuits that mediate other behaviors. The neurotransmitter composition of 3Ns varied less across modalities than that of 2Ns, but we again observed differences between the excitatory/inhibitory balance of SEZ and non-SEZ outputs that may reflect how taste input is used to regulate feeding circuits in the SEZ compared to circuits for other behaviors outside the SEZ.

### Connecting taste pathways to behavior

This study provides a global view of early taste processing in the fly brain. We did not characterize taste circuits beyond 3Ns because iterative tracing generates larger and larger numbers of neurons at each layer, and within a few steps one would reach a large proportion of neurons in the entire brain.^49^ In our whole-brain simulations, we found that stimulation of GRNs activated a large proportion of 2Ns but only a minority of 3Ns. This is partially due to 3Ns that are inhibited rather than excited by taste input, but it is also likely that some 3Ns receive taste input that is too weak to elicit neuronal firing on its own and instead contributes to neuronal firing in combination with other stimuli. Thus, while the number of neurons receiving taste input clearly expands at each layer, the proportion of neurons that receive strong excitatory input may decrease, potentially reflecting a transformation from neurons dedicated to taste processing to neurons with integrative roles.

Recent studies investigating sugar-evoked locomotor stopping and proboscis extension suggest that sensory processing of taste information likely occurs within the first few layers of the circuit, with downstream neurons (fourth-order neurons and beyond) comprising premotor and motor pathways.^25,39,40^ Thus, the 2Ns and 3Ns identified here may represent the core neurons that perform computations related to sensory processing before downstream circuits implement sensory-motor transformations. The taste neurons identified in this study represent candidate neurons whose functional roles can be tested in future work, given the availability of genetic tools to target individual cell types in the SEZ and other regions.^6–11^ It will be interesting to compare the taste coding and behavioral roles of neurons belonging to different 2N and 3N classes that we identified, such as excitatory versus inhibitory neurons, SEZ versus non-SEZ neurons, and neurons projecting to different brain regions.

### Limitations of the study

This study relied on connectomic analyses to trace taste pathways; functional studies will be needed to confirm the role of specific neurons in taste processing. In addition, the connectome represents the brain of a single female fly at a single moment in time, and it is possible that neuronal connectivity varies across individuals, sexes, or a fly’s lifetime. Finally, we focused on tracing taste inputs from the labellum, and future work will be needed to identify pathways that process taste input from other organs.

## ACKNOWLEDGMENTS

We thank Chris Rodgers for input on the code and figures, members of the Devineni lab for comments and feedback, and Philip Shiu for publicly sharing code for brain simulations. We acknowledge the Princeton FlyWire team and members of the Murthy and Seung labs for development and maintenance of FlyWire (supported by BRAIN Initiative grant MH117815 to Murthy and Seung). This work was supported by the NIH (R01DC021478 to A.V.D.) and the Whitehall Foundation (grant 2022-08-017 to A.V.D.).

## AUTHOR CONTRIBUTIONS

A.V.D. conceived and supervised the project, performed model simulations, and edited the analysis code. S.R.W. and M.P-G. wrote initial drafts of the analysis code, performed analyses, and generated drafts of the figures. A.V.D. wrote the manuscript and finalized the figures with input from all authors.

## STAR METHODS

### RESOURCE AVAILABILITY

#### Lead contact

Further information and requests for resources and reagents should be directed to and will be fulfilled by the lead contact, Anita Devineni.

#### Materials availability

This study did not generate new unique reagents.

#### Data and code availability

- All data reported in this paper can be obtained using the publicly available code shared on GitHub (see below). Data files containing lists of second- and third-order neurons identified in this study are also available in the same GitHub repository.
- Code and associated datasets needed to perform all analyses in this study are publicly available on GitHub at https://github.com/avdevineni/taste-connectome.
- Any additional information required to reanalyze the data reported in this paper is available from the lead contact upon request.

## METHOD DETAILS

### Identifying GRNs, 2Ns, and 3Ns

Datasets containing information about neurons, connectivity, and synapse locations were downloaded from FlyWire at https://codex.flywire.ai/api/download. All analyses in this study used version 630. Analyses were performed in Python, and the code and associated datasets needed to perform all analyses in this study are available on Github (https://github.com/avdevineni/taste-connectome).

We focused on tracing the outputs of GRNs in the left hemisphere (referred to in some previous papers^24,25^ as the right hemisphere due to the left-right inversion of the brain volume that was discovered later^4,5^), which have been more thoroughly annotated. GRNs for each modality were identified based on lists from previous studies,^17,18^ neuron annotations in FlyWire, and manual inspection of each neuron’s morphology. Our GRN lists are very similar to those used in Shiu et al.,^17^ with the following exceptions: we excluded one sugar GRN whose identity as a sugar versus water GRN seemed uncertain, we excluded one bitter GRN that has no connections and is therefore irrelevant for analysis, we excluded 7 IR94e GRNs that have a different morphology from other IR94e GRNs and were not included in IR94e tracing analyses by Guillemin et al.,^18^ and we added one sugar GRN that has been annotated as such in FlyWire and appears to have the correct morphology.

We used a connection threshold of 5 synapses to identify 2Ns, which is the threshold suggested by FlyWire to exclude false positives.^4^ We set a more stringent threshold of 10 synapses to identify 3Ns (including 10 synapses for both GRN-2N and 2N-3N connections), which reduced the number of 3Ns by ∼50% compared to a 5-synapse threshold. Analyses of connection stereotypy across hemispheres and brains suggest that a value of ∼10 synapses represents the threshold for reliable connections.^5^ Thus, it is likely that we have identified all reliable 3Ns, whereas some of the 2Ns with weaker GRN input (< 10 synapses) may not be reliably connected to GRNs in all flies. Both 2N and 3N lists excluded GRNs of all modalities; lists of 3Ns excluded neurons that were also 2Ns for the same modality, but not other modalities. The 2N-3N connectivity heatmap (Figure 3F) uses the same criteria as used for 3N identification, meaning that 2N-2N connections within the same modality are not shown.

### Analyzing neuronal outputs

Depending on the analysis, we focused on analyzing either output synapses or output connections, with a “connection” comprising all synapses between a single pair of presynaptic and postsynaptic neurons in a given neuropil and requiring a 5-synapse threshold. The term “output” may refer to either an output synapse or output connection, depending on the context, and figure legends specify which metric is being quantified. Analyses of 2N outputs (e.g., analyzing neurotransmitter types or neuropil location), did not exclude outputs to GRNs or other 2Ns; similarly, analyses of 3N outputs did not exclude outputs to GRNs, 2Ns, or other 3Ns. Methods for specific analyses are described further below.

### Analyzing neurotransmitter types and excitatory/inhibitory convergence

For analysis of excitatory and inhibitory neurotransmission, we focused on synapses predicted to use acetylcholine, GABA, or glutamate, as it is not clear whether other predicted neurotransmitters (dopamine and serotonin) would have an excitatory or inhibitory effect on downstream cells. Only a very small fraction of synapses were predicted to use these other neurotransmitters. The FlyWire datasets include neurotransmitter predictions at both the synapse level and the neuron level, with the neuron predictions derived by pooling the synapse predictions and assuming that each neuron only expresses one neurotransmitter according to Dale’s principle.^4^ When quantifying the neurotransmitter types for output synapses or output connections, we used synapse-level predictions. When analyzing excitatory/inhibitory convergence onto 3Ns (Figure 4A-B), we used neuron-level predictions because it was essential to count each 2N-3N pair only once and the synapse-level predictions for the neurotransmitter type of 2N-3N connections sometimes conflicted, even for connections between the same 2N-3N pair. Plots of net excitation onto 3Ns (Figure 4C) also used neuron-level predictions.

Analyses of excitatory/inhibitory convergence onto 3Ns (Figure 4A-B) focused on 3Ns receiving exactly two or three 2N inputs, which comprise 86% of all 3Ns receiving more than one 2N input. We excluded cases where at least one input was predicted to use a neurotransmitter other than acetylcholine, GABA, or glutamate. For 3Ns receiving exactly two 2N inputs, our dataset included 71-112 3Ns (112 for sugar, 71 for water, 98 for bitter, and 72 for IR94e), representing 50-97% of all 3Ns receiving more than one 2N input (50% for sugar, 76% for water, 91% for bitter, 97% for IR94e). For 3Ns receiving exactly three 2N inputs, our dataset included 2-53 3Ns (53 for sugar, 12 for water, 8 for bitter, and 2 for IR94e), representing 3-24% of all 3Ns receiving more than one 2N input (24% for sugar, 13% for water, 7% for bitter, 3% for IR94e).

Because small numbers of neurons do not provide reliable data, when presenting results for 3Ns with three 2N inputs we only included data with at least 10 neurons (sugar and water), and we note that the water data (12 neurons) should be interpreted cautiously. The expected proportion of 3Ns in each category (all excitatory inputs, all inhibitory inputs, or a combination of excitatory and inhibitory inputs) was calculated using probability rules based on the proportion of excitatory and inhibitory 2N inputs onto the specific 3Ns being analyzed (note that these proportions may differ from those when considering all 2N inputs onto all 3Ns). For example, if the proportions of excitatory and inhibitory 2N inputs are 0.3 and 0.7, respectively, then the expected proportion of 3Ns receiving two excitatory inputs, two inhibitory inputs, or one input of each type is 0.3*0.3, 0.7*0.7, or 2*0.3*0.7, respectively.

### Analyzing projection targets of 2Ns and 3Ns

To classify neuronal outputs within versus outside of the SEZ, we defined the SEZ as encompassing the following regions: gnathal ganglia (GNG), prow (PRW), saddle (SAD), flange (FLA), and cantle (CAN). This definition includes periesophageal neuropils except for the antennal mechanosensory and motor center, which has sometimes been categorized as an SEZ neuropil^9^ but is clearly separated and has specific functions (e.g., auditory processing^50,51^) that are distinct from SEZ processing. To be classified as non-local neurons, 2Ns or 3Ns needed to have at least one output connection outside the SEZ (with a 5-synapse threshold). Classification of laterality for non-SEZ output synapses (Figures 2F and 5G) was based on whether they were located in the left or right hemisphere for brain regions that are lateralized (which includes the majority of brain regions outside the SEZ); we excluded synapses in regions that are not lateralized. When analyzing non-SEZ brain regions receiving 2N or 3N outputs (Figures 2G and 5H), we pooled outputs located in homologous regions of both hemispheres. Plots of SEZ versus non-SEZ outputs (Figures 2C and 5D), ipsilateral versus contralateral outputs (Figures 2F and 5G), and outputs in different brain regions (Figure 2G and 5H) rely on synapse counts, whereas plots categorizing the neurotransmitter types of neuronal outputs (Figures 2ṣD-E and 5E-F) rely on the number of output connections.

For plots of brain regions containing 2N or 3N outputs outside the SEZ (Figures 2G and 5H), we included the regions containing the most numerous output synapses for each modality. For 2Ns, we included the top 6 regions for each modality, comprising a total of 10 regions that included >96% of non-SEZ output synapses for each modality. For 3Ns, we included the top 12 regions for each modality, comprising a total of 19 regions that included >97% of non-SEZ output synapses for sugar, bitter, and IR94e 3Ns and 91% for water 3Ns.

Brain region abbreviations not defined in the main text include: superior intermediate protocerebrum (SIP); inferior clamp (ICL), crepine (CRE), inferior bridge (IB), antennal lobe (AL), mushroom body calyx (MB_CA), mushroom body vertical lobe (MB_VL), posterior ventrolateral protocerebrum (PVLP), posterior lateral protocerebrum (PLP), wedge (WED), gorget (GOR).

In the figures, brain regions are grouped by location or function: superior protocerebrum (SMP, SIP, SLP), inferior neuropils (SCL, ICL, CRE, IB), mushroom body (MB_CA, MB_VL), olfactory regions (AL, LH), ventrolateral neuropils (AVLP, PVLP, PLP, WED), ventromedial neuropils (VES, GOR, SPS, IPS), and lateral complex (LAL).

### Whole-brain simulations

Brain simulations were conducted in Python based on code provided publicly by Philip Shiu (https://github.com/philshiu/Drosophila_brain_model), and details about the model are described in Shiu et al. (2023).^17^ We used the default parameters for the model, which were chosen by Shiu et al.^17^ based on published experimental data and include: -52 mV resting potential, -52 mV reset potential after a spike, -45 mV spiking threshold, 2.2 ms refractory period, 5 ms tau for synapse decay, 1.8 ms time delay from spike to change in membrane potential, and 0.275 mV synaptic weight. In all simulations, neurons were stimulated for 1 second. All simulations were repeated over 30 trials, and mean values were quantified. We did not apply a threshold for activation; all neurons that had an average firing rate above zero were considered to be “activated” in a given simulation.

## QUANTIFICATION AND STATISTICAL ANALYSIS

Statistical tests are described in the figure legends. Ordinary least squares regression was implemented in Python to obtain r-squared values and p-values for correlations in Figures 1G and 3I. Other statistical tests were performed using GraphPad Prism, Version 9, including Fisher’s exact test to compare proportions between different modalities in Figures 2B-F and 5C-G, and chi-squared test to compared observed versus expected values in Figure 4A-B.

